# A New Platform for Label-Free, Proximal Cellular Pharmacodynamic Assays: Identification of Glutaminase Inhibitors Using Infrared Matrix-Assisted Laser Desorption Electrospray Ionization Mass Spectrometry

**DOI:** 10.1101/2023.01.30.526319

**Authors:** Fan Pu, Andrew J. Radosevich, Brett G. Bruckner, David A. Fontaine, Sanjay C. Panchal, Jon D. Williams, Sujatha M. Gopalakrishnan, Nathaniel L. Elsen

**Affiliations:** Discovery Research, AbbVie Inc., 1 North Waukegan Rd., North Chicago, IL, USA 60064

**Keywords:** high throughput, glutaminase, cellular assay, biochemical assay, mass spectrometry

## Abstract

Cellular pharmacodynamic assays are crucial aspects of lead optimization programs in drug discovery. These assays are sometimes difficult to develop, oftentimes distal from the target and frequently low throughput which necessitates their incorporation in the drug discovery funnel later than desired. The earlier direct pharmacodynamic modulation of a target can be established, the less resources are wasted on compounds that are acting via an off-target mechanism. Mass spectrometry is a versatile tool that is often used for direct, proximal cellular pharmacodynamic assay analysis but liquid chromatography-mass spectrometry methods are low throughput and unable to fully support structure-activity relationships efforts in early medicinal chemistry programs. Infrared matrix-assisted laser desorption electrospray ionization (IR-MALDESI) is an ambient ionization method amenable to high throughput cellular assays, capable of diverse analyte detection, ambient and rapid laser sampling process, and low cross contamination. Here we demonstrate the capability of IR-MALDESI for detection of diverse analytes directly from cells and report the development of a high throughput label free, proximal cellular pharmacodynamic assay using IR-MALDESI for discovery of glutaminase inhibitors and a biochemical assay for hit confirmation. We demonstrate the throughput with a ∼100,000 compound cellular screen. Hits from the screening were confirmed by retesting in dose-response with mass spectrometry-based cellular and biochemical assays. A similar workflow can be applied to other targets with minimal modifications, which will speed up discovery of cell active lead series and minimize wasted chemistry resources on off-target mechanisms.

## Introduction

Cellular pharmacodynamic (PD) assays measure drug-induced cellular responses and are crucial for lead optimization programs in drug discovery. These assays can be difficult to develop and low throughput, which necessitates their incorporation in the drug discovery funnel later than desired. In addition, they oftentimes detect a PD event distal from the target thereby increasing the chance that an off-target mechanism is responsible for the observed biological response. The earlier direct PD modulation of a target can be established, the less resources are wasted on compounds that are inactive in cells or active by off-target mechanisms. Mass spectrometry (MS) is an analytical tool with superior specificity that can be used to directly measure intracellular metabolites proximal to the target, making it an excellent readout for cellular PD assays. However, the requirement of chromatography separation and sometimes intensive sample preparations has severely limited the throughput.

Use of MS without chromatography can dramatically increase throughput while retaining the advantages of a MS-based assay.^1^ The tradeoff is typically loss of sensitivity and increased variability. However, there are still numerous applications in the early drug discovery phase that could benefit from a high throughput platform with sufficient analytical performance. The two most well-known methods which implement this strategy are RapidFire (with or without solid phase extraction)^2^ and matrix-assisted laser desorption/ionization (MALDI)^3^. More novel ionization methods reported in recent years have enabled even higher throughput that match or even exceed the speed of conventional optical-based methods. Examples of such technologies include acoustic mist ionization (AMI)^4^, Acoustic droplet ejection-open port interface (ADE-OPI)^5^, desorption electrospray ionization (DESI)^6^, and liquid atmospheric pressure-MALDI^7^. The applications of these methods were initially concentrated on biochemical assays. Recently, applications to cellular assays are gaining interest, some examples are the use of AMI for high throughput metabolomics profiling^8^ and a mechanistic MALDI cellular assay based on measurement of intracellular metabolite build up^9^.

Infrared matrix-assisted laser desorption electrospray ionization (IR-MALDESI) is an ambient ionization method that ionizes molecules with electrospray post laser ablation.^10, 11^ The optimal matrix for IR-MALDESI is water, making it amenable to biological samples. We have constructed a high-throughput IR-MALDESI system in house and have reported its use for biochemical assays^12^ and intact protein analysis^13^. A scanning speed of up to 22 samples per second is achievable although 2 samples per second is more commonly used in assays^14^. IR-MALDESI has also been reported for MS imaging applications where a variety of analytes including metabolites and lipids can be covered^15, 16^, hence it should be applicable to high throughput cellular assays with appropriate assay protocols.

Glutaminase is an enzyme that catalyzes the conversion of glutamine (Gln) to glutamate (Glu), which then feeds into the tricarboxylic acid cycle. Gln is a key amino acid that fuels the growth of many cancer cells, and it has been established that targeting Gln utilization by developing glutaminase inhibitors is a viable strategy for cancer treatment.^17^ CB839 (telaglenastat)^18^ is the only glutaminase inhibitor that has advanced into clinical trials, although development was recently discontinued due to lack of clinical efficacy.

Here we show that a high throughput IR-MALDESI workflow can provide coverage for a wide range of cellular metabolites and lipids directly from cells and report the development of a high throughput cellular PD assay and an ancillary biochemical assay to identify glutaminase inhibitors using IR-MALDESI. We screened ∼100,000 compounds in the cellular glutaminase PD assay, and performed hit confirmation with the biochemical assay. IR-MALDESI assays were performed at a speed of 3 minutes per 384-well plate (2 samples per second). CB839 was in the screening library and was identified and confirmed by the IR-MALDESI assays.

## Results and Discussion

### Cellular assay development

To take full advantage of the ability of IR-MALDESI to sample from any plate type we set out to design an assay workflow that minimized the use of consumable plastics from start to finish. To do so we first dispensed compounds to empty plates using an Echo, then dispensed cells to the compound containing plates. This workflow works for suspension cell lines, but for adherent cell lines cells need sufficient time to attach in the assay time window before washing. A549 cells can attach to assay plate in as soon as one hour, as evidenced by a similar observed confluency before and after cell wash (data not shown). The assay protocol for an IR-MALDESI cell-based assay is illustrated in Figure 1A. Gln was present in the cell culture media; therefore, cells were washed twice with MS friendly buffer (150 mM ammonium acetate) before lysing the cells to analyze intracellular metabolites. This wash step can be reduced or omitted for some assays where analytes of interest would not be interfered with by cell culture media components. For cell lysis and extraction, we used a mixture of 20:80 methanol/water (v/v). Water, greater than 50% v/v, needs to be included for optimal IR laser ablation. It is also possible to perform cell lysis and extraction with pure organic solvent, then add water or reconstitute in high water content solvent. The high throughput IR-MALDESI system we constructed is compatible with any standard microtiter plate, thus all steps including MS readout can be performed in the same cell culture plate. This workflow resulted in a cell-based assay which uses a single 384 well plate and zero pipet tips. Total consumable plastics cost for the assay was 1.3 cents/well.

**Figure 1.**
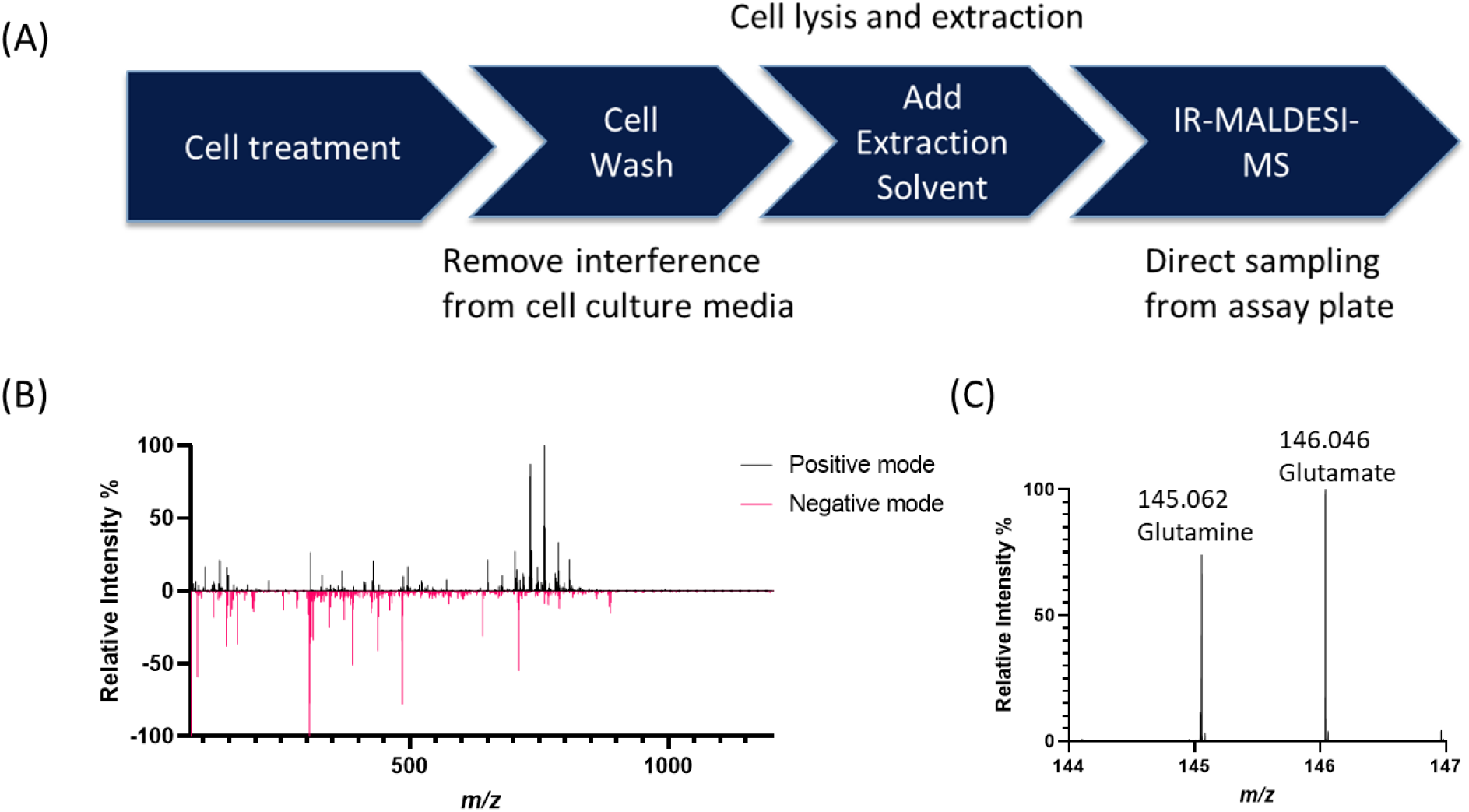
Development of cellular assay. (A) Assay workflow for IR-MALDESI cellular assay; (B) example mass spectrum showing rich features obtained from A549 cells using IR-MALDESI, spectra of each polarity were background subtracted then normalized in their respective range (m/z 75-300 and 300-1200) before combined. (C) A mass spectrum zoomed-in on Gln and Glu, the peaks can be clearly differentiated.

To examine the effectiveness and analyte coverage by this IR-MALDESI workflow, we performed a cell titration experiment using A549 cells. A high resolving power of 240,000 (FWHM at *m/z* = 200) was used for this experiment. A combined mass spectrum obtained using IR-MALDESI is shown in Figure 1B. Putative annotation was performed using METASPACE with an FDR less than 10 and 3 ppm *m/z* tolerance. By checking the titration pattern of intensity heatmaps, we were able to separate cellular features from background (Figure S1). When combining results from both polarities, a total of 404 unique analytes matching the titration pattern were found and annotated. Detailed annotation results are provided in Supplementary xlsx file.

By setting a smaller *m/z* range, it is easy to detect Gln and Glu and their peaks are clearly differentiated (Figure 1C). To find the optimal assay time, we tested dose response properties using CB839. As is shown in Figure S2, the assay window widens with longer incubation time and stabilizes after 4 hours, although IC_50_ values were similar regardless of assay time. Therefore, 4 hours incubation was selected as the screening condition.

### Biochemical assay development

One major advantage of the MS-based biochemical assays is the capability to measure substrate and product of the enzymatic reaction directly and simultaneously without labeling or coupling reactions. This simplifies assay development. Glutaminase enzymatically catalyzes the conversion of Gln to Glu (Figure 2A), both of which can be readily detected by IR-MALDESI in buffered solution with detection limits of at least 15 μM for Gln (noise was nearly zero) and around 30 μM for Glu (signal-to-noise ratio > 3), respectively (Figure S3). Glutaminase C (GAC) is an isoform of glutaminase, and it was used for the biochemical assay.

**Figure 2.**
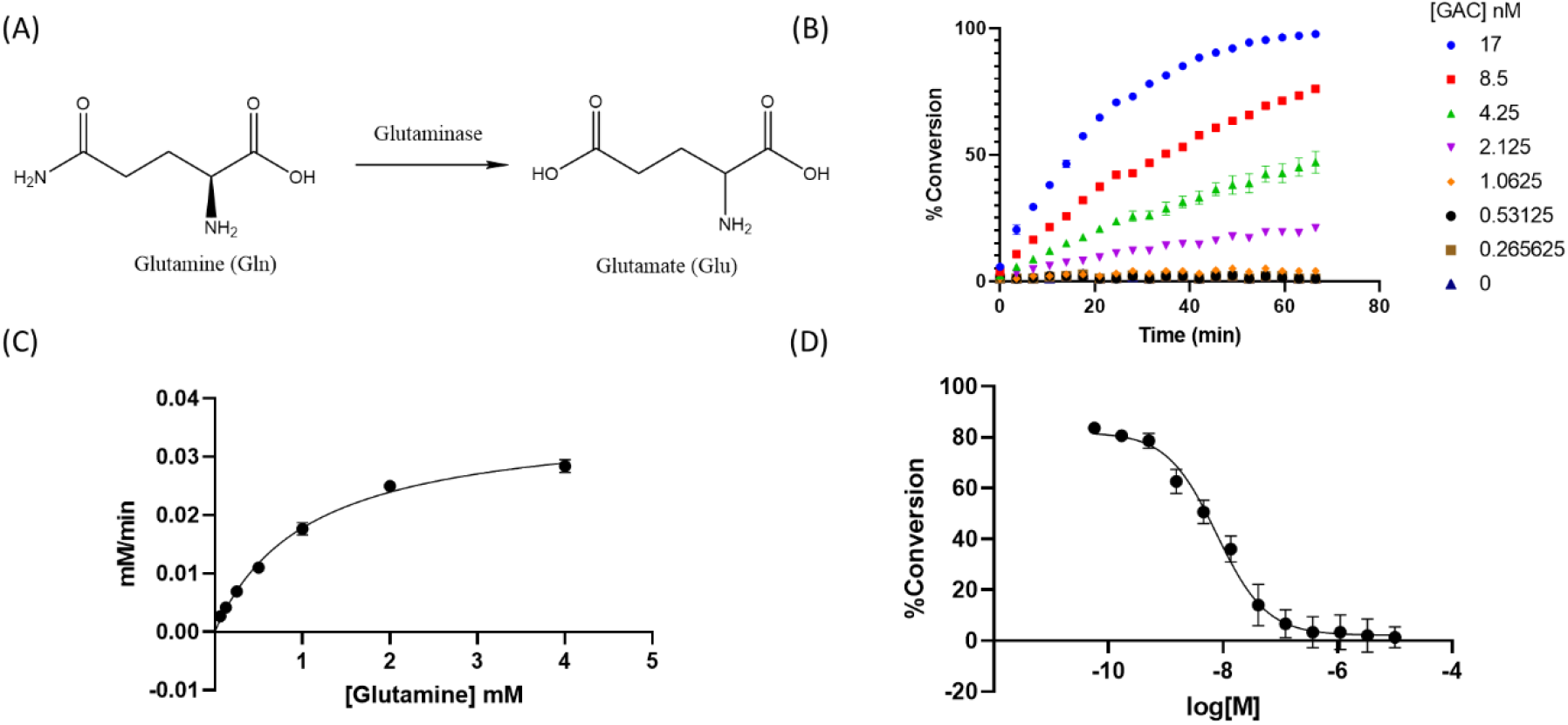
Development of biochemical assay. (A) Glutaminase convert glutamine to glutamate; (B) enzyme titration of GAC; (C) K_m_ curve obtained from titration of glutamine; (D) dose response curve of tool compound CB839.

Titration of GAC enzyme was performed with 2 mM glutamine to determine suitable enzyme concentration and reaction time. A 1:2 serial titration of GAC starting from 17 nM was carried out in triplicate in a 384-well Proxiplate. IR-MALDESI was operated in kinetic mode where the same samples were analyzed every 3 min without quenching. The results are presented in Figure 2B, 4 nM GAC and 45 minutes reaction were selected for the assay to ensure a large assay window while still operating in the linear region.

A 1:2 serial titration of substrate Gln was then performed with 4 nM GAC, with highest Gln concentration being 4 mM. Similarly, IR-MALDESI was operated in kinetics mode and all reaction progression data were fitted by linear fitting (Figure S4). The slopes were plotted against Gln to obtain a K_m_ value (Figure 2C) by fitting to the Michaelis-Menten model. The K_m_ for GAC was determined to be 1.05 mM, which is comparable to a reported value of 2.1 mM in the presence of 50 mM phosphate^19^. 2 mM Gln was chosen for the biochemical assay.

With the assay conditions established, we tested the biochemical assay with tool compound CB839. An IC_50_ curve was obtained by 1:3 serial titration of CB839 from 10 μM in triplicate (Figure 2D). The IC_50_ was determined to be 7.7 nM, similar to literature reported value of <50 nM^18^. Using 10 μM CB839 as positive control and DMSO as negative control, robust Z’ above 0.9 can be reliably achieved.

### Pilot screen and screening funnel

A pilot screen of 1080 well-annotated compounds containing FDA approved compounds and bioactive compounds from Selleckchem was conducted at 60 μM using the cellular assay. Each compound plate was tested in duplicate. As shown in Figure 3A, the two replicates showed good correlation. Robust Z’ of 0.79 ± 0.11 was achieved from the 6 pilot screen plates. Hits were defined as compounds with percent inhibition above 50%. CB839 was in the pilot screen and was identified as a hit. The other two hit compounds were crystal violet and sanguinarine chloride. The hit rate of the pilot screen was 0.3% and the assay performance was deemed suitable for a larger scale screen.

We devised our screening funnel to consist of cost-effective IR-MALDESI based assays (Figure 3B). Primary screening was conducted using the cellular PD assay, hits from the screen were confirmed with both cellular PD and biochemical assays. Hits confirmed in both assays can be further characterized by biophysical methods, which is out of the scope of this work.

**Figure 1.**
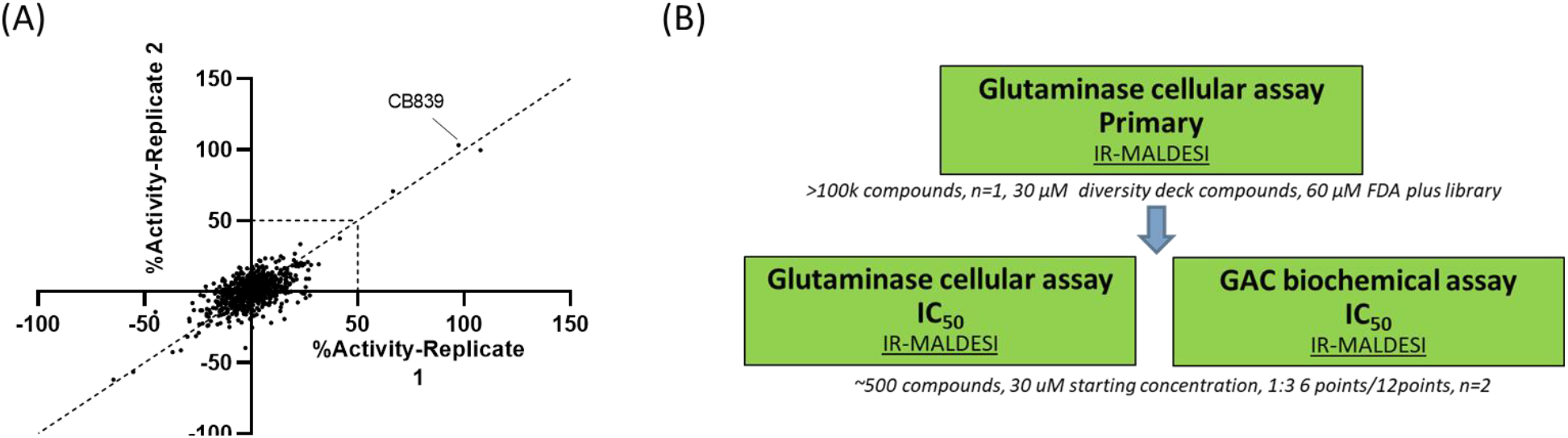
(A) Pilot screen showed good correlation between two replicates. (B) Screening funnel that consisted of IR-MALDESI assays with high throughput cellular assay being the primary screen method.

### Library screen and hit confirmation

For the primary screen, we screened the entire FDA plus library (2494 compounds tested representing FDA approved compounds and other bioactive compounds) at 60 μM, and another ca. 102 000 compounds from the main screening library at 30 μM. Robust Z’ of the 293 screen plates was 0.68 ± 0.19. Like the pilot screen, compounds with higher than 50% inhibition were considered hits and a total of 561 compounds were picked according to this criterion, a 0.5% pick rate. Figure 4A shows a scatter plot of the single point screening results of ∼100,000 compounds.

**Figure 4.**
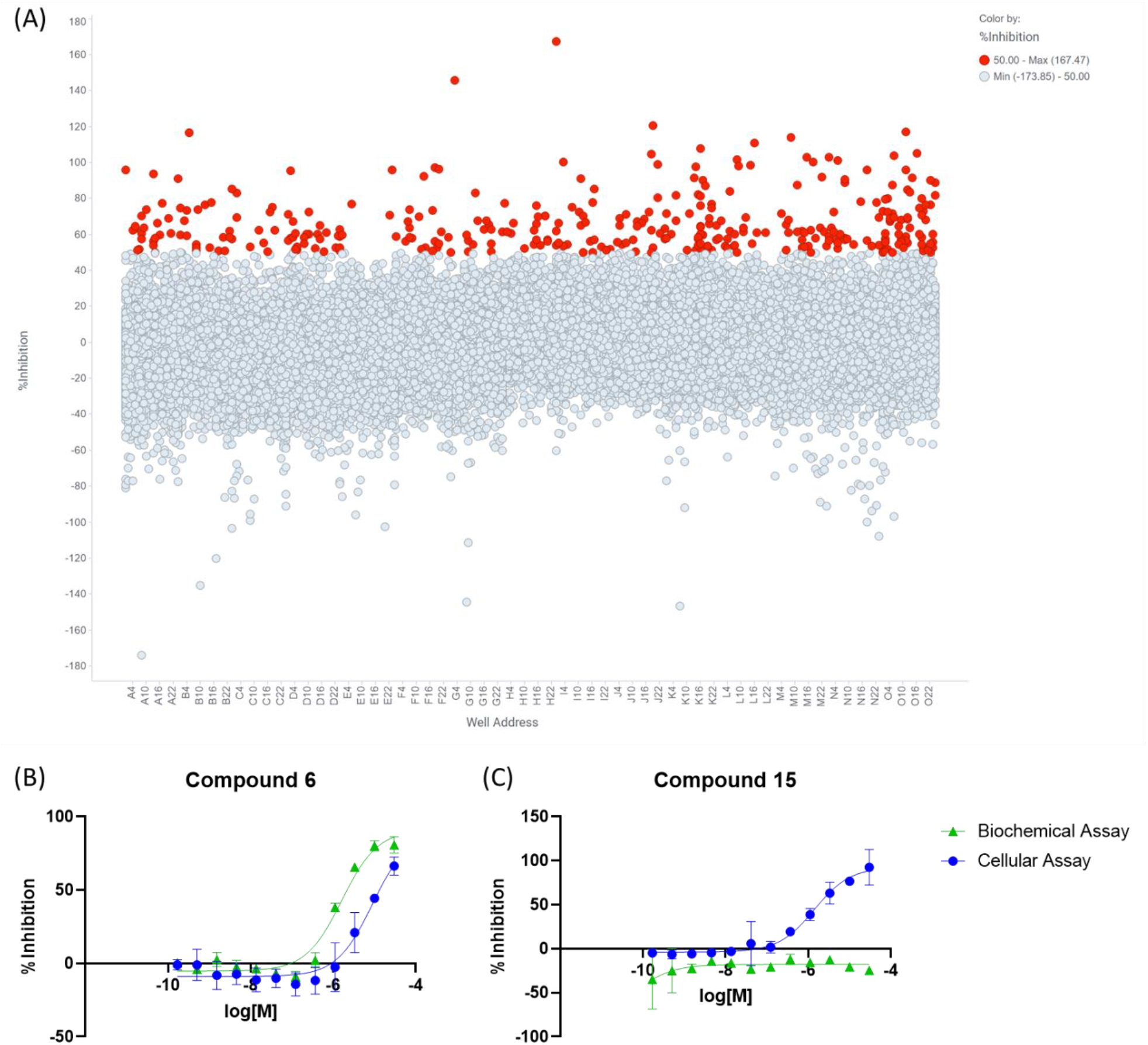
(A) Scatter plot of primary screen results. Hits (>50% inhibition) are labeled red. Dose response curves of (B) compound 6 (active in both assays) and (C) compound 15 (active only in cellular assay).

Picks from the screen were tested in dose response in both biochemical and cellular assays using IR-MALDESI. 22 compounds were confirmed active with suitable curve shapes in the cellular assay. Among the confirmed hits, 15 also showed inhibition in the biochemical assay; 7 compounds were only active in the cellular assay, indicating they are likely active through off-target mechanism or inhibiting other glutaminase isoforms. Examples of dose response curves for each case are shown in Figure 4B and C and the rest of the hit compounds are shown in Table S1.

## Conclusions

We have shown that direct cellular detection by IR-MALDESI can analyze a wide variety of intracellular metabolites and lipids and developed a MS-based cellular PD assay that directly measures intracellular conversion from Gln to Glu. Most notably, the IR-MALDESI readout speed on a cell plate is the same as a biochemical assay (3 minutes per 384-well plate or 2 samples per second), which makes it possible to be utilized earlier in the screening funnel to accelerate drug discovery. We demonstrated this with an extreme scenario where our primary screen was the cellular PD assay then used an IR-MALDESI biochemical assay and the cellular assay for hit confirmation. ∼100 000 compounds were screened with good robust Z’. ∼500 hits were identified and 22 were confirmed in the cellular assay. Importantly, CB839, a known glutaminase inhibitor, was found in the screen and confirmed active in both cellular and biochemical assays. A full deck screen with millions of compounds tested might lead to more chemically attractive confirmed hits that bind to the target. Screening with cellular PD assay can provide cell active starting points, identify pathway inhibitors and additional mechanisms. It is also easy to start a HTS campaign if the recombinant protein is not readily available. The downside might be that cell impermeable compounds will be missed. This novel high throughput technology can provide label free proximal cellular PD assays for many drug targets, including some that are not possible by traditional high throughput optical assays, and has the potential to reduce costs and discovery timelines.

## Methods

### Details on Materials, IR-MALDESI and MS, Cell Culture and Compound Dispensing are provided in supporting information

#### Glutaminase Cellular Assay

Cells were detached from flasks using 0.25% Trypsin-EDTA (Gibco) and counted using Vi-cell XR cell viability analyzer (Beckman Coulter). The cells were then diluted with cell culture media to a concentration of *ca*. 500,000 cells per mL and dispensed to assay plates at 20 μL per well using a Multidrop Combi (Thermo Fisher). Final cell density was 10,000 cells per well. The compound screening concentration was 30 μM under these experimental conditions (60 μM for FDA plus library). The assay plates were then incubated at 37 °C, 5% CO_2_ for 4 hours, which allowed the cells to attach and compound effect to accumulate. After incubation, the assay plates were washed twice with 150 mM ammonium acetate using a Blue®Washer (BlueCatBio). 20:80 methanol/water (v/v) was dispensed to the washed plates at 100 μL per well to lyse cells and extract analytes. The assay plates were directly analyzed with IR-MALDESI without any transfer.

#### GAC Biochemical Assay

Assay buffer for GAC enzymatic assay was 50 mM potassium phosphate buffer (pH 7.4) with 0.01% Tween 20. Buffered enzyme 2X solution (8 nM GAC) was first dispensed to assay plates at 2.5 μL per well and pre-incubated with compounds for 15 min. 2.5 μL of buffered substrate 2X solution (4 mM glutamine) was then dispensed to assay plates to initiate the reaction. After 45 min, the reaction was quenched by adding 15 μL per well of 0.1% formic acid. The assay plates were directly analyzed with IR-MALDESI without any transfer.

#### Data Analysis

Annotation of metabolites and lipids was performed by converting raw data first to mzML file using MSConvert from ProteoWizard, then convert to imzML file using imzMLConverter. imzML files were uploaded to METASPACE annotation platform (https://metaspace2020.eu/) for annotation.^20^ An FDR of less than 10 and *m/z* tolerance of 3 ppm were used to search HMDB v4 and LipidMaps (2017-12-12) databases.

An in-house data browser was used to process all IR-MALDESI targeted assay data. Peak intensities of *m/z* 145.062 (Gln) and 146.046 (Glu) were extracted from raw data files with a mass tolerance of 5 ppm. Triplicate spectra collected from the same well were averaged and mapped to their well location. Percent conversion was calculated using the following equation:

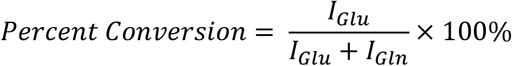

I_Glu_ and I_Gln_ were averaged peak intensity of Glu and Gln, respectively.

Robust Z’ was calculated with robust standard deviation (rSD) and median using the following equation:

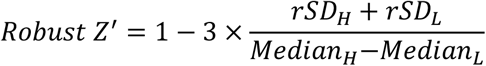

Subscript H and L stands for high control (DMSO) and low control (CB839 treatment). rSD was calculated as:

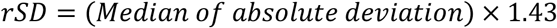

After data processing with data browser, single text files containing a list of well address and percent conversion for each assay plate were generated and loaded to in-house HTS software to link compound identity to MS data. The data were then visualized using TIBCO Spotfire 10 and exported for making graphs. All graphs were made using GraphPad Prism 9 or Microsoft PowerPoint except for Figure 4A, which was exported from TIBCO Spotfire 10.

## Supporting information

Annotation list

Supportin Information

## Acknowledgements

F.P., A.R., B.B., S.P., J.W., S.G. and N.E. are employees of AbbVie. D.F. was an employee of AbbVie at the time of the study. The design, study conduct, and financial support for this research were provided by AbbVie. AbbVie participated in the interpretation of data, review, and approval of the publication.

## References

1. Pu, F.; Elsen, N. L.; Williams, J. D., Emerging Chromatography-Free High-Throughput Mass Spectrometry Technologies for Generating Hits and Leads. ACS Med Chem Lett 2020, 11 (11), 2108–2113.

2. Bretschneider, T.; Ozbal, C.; Holstein, M.; Winter, M.; Buettner, F. H.; Thamm, S.; Bischoff, D.; Luippold, A. H., RapidFire BLAZE-Mode Is Boosting ESI-MS Toward High-Throughput-Screening. SLAS Technol 2019, 24 (4), 386–393.

3. Haslam, C.; Hellicar, J.; Dunn, A.; Fuetterer, A.; Hardy, N.; Marshall, P.; Paape, R.; Pemberton, M.; Resemannand, A.; Leveridge, M., The Evolution of MALDI-TOF Mass Spectrometry toward Ultra-High-Throughput Screening: 1536-Well Format and Beyond. J Biomol Screen 2016, 21 (2), 176–86.

4. Sinclair, I.; Bachman, M.; Addison, D.; Rohman, M.; Murray, D. C.; Davies, G.; Mouchet, E.; Tonge, M. E.; Stearns, R. G.; Ghislain, L.; Datwani, S. S.; Majlof, L.; Hall, E.; Jones, G. R.; Hoyes, E.; Olechno, J.; Ellson, R. N.; Barran, P. E.; Pringle, S. D.; Morris, M. R.; Wingfield, J., Acoustic Mist Ionization Platform for Direct and Contactless Ultrahigh-Throughput Mass Spectrometry Analysis of Liquid Samples. Anal Chem 2019, 91 (6), 3790–3794.

5. Zhang, H.; Liu, C.; Hua, W.; Ghislain, L. P.; Liu, J.; Aschenbrenner, L.; Noell, S.; Dirico, K. J.; Lanyon, L. F.; Steppan, C. M.; West, M.; Arnold, D. W.; Covey, T. R.; Datwani, S. S.; Troutman, M. D., Acoustic Ejection Mass Spectrometry for High-Throughput Analysis. Anal Chem 2021, 93 (31), 10850–10861.

6. Morato, N. M.; Holden, D. T.; Cooks, R. G., High-Throughput Label-Free Enzymatic Assays Using Desorption Electrospray-Ionization Mass Spectrometry. Angewandte Chemie 2020, 132 (46), 20639–20644.

7. Krenkel, H.; Brown, J.; Richardson, K.; Hoyes, E.; Morris, M.; Cramer, R., Ultrahigh-Throughput Sample Analysis Using Liquid Atmospheric Pressure Matrix-Assisted Laser Desorption/Ionization Mass Spectrometry. Anal Chem 2022, 94 (10), 4141–4145.

8. Smith, M. J.; Ivanov, D. P.; Weber, R. J. M.; Wingfield, J.; Viant, M. R., Acoustic Mist Ionization Mass Spectrometry for Ultrahigh-Throughput Metabolomics Screening. Anal Chem 2021, 93 (26), 9258–9266.

9. Weigt, D.; Parrish, C. A.; Krueger, J. A.; Oleykowski, C. A.; Rendina, A. R.; Hopf, C., Mechanistic MALDI-TOF Cell-Based Assay for the Discovery of Potent and Specific Fatty Acid Synthase Inhibitors. Cell Chem Biol 2019, 26 (9), 1322–1331 e4.

10. Sampson, J. S.; Murray, K. K.; Muddiman, D. C., Intact and top-down characterization of biomolecules and direct analysis using infrared matrix-assisted laser desorption electrospray ionization coupled to FT-ICR mass spectrometry. J Am Soc Mass Spectrom 2009, 20 (4), 667–73.

11. Caleb Bagley, M.; Garrard, K. P.; Muddiman, D. C., The development and application of matrix assisted laser desorption electrospray ionization: The teenage years. Mass Spectrom Rev 2021.

12. Pu, F.; Radosevich, A. J.; Sawicki, J. W.; Chang-Yen, D.; Talaty, N. N.; Gopalakrishnan, S. M.; Williams, J. D.; Elsen, N. L., High-Throughput Label-Free Biochemical Assays Using Infrared Matrix-Assisted Desorption Electrospray Ionization Mass Spectrometry. Anal Chem 2021, 93 (17), 6792–6800.

13. Pu, F.; Ugrin, S. A.; Radosevich, A. J.; Chang-Yen, D.; Sawicki, J. W.; Talaty, N. N.; Elsen, N. L.; Williams, J. D., High-Throughput Intact Protein Analysis for Drug Discovery Using Infrared Matrix-Assisted Laser Desorption Electrospray Ionization Mass Spectrometry. Anal Chem 2022, 94 (39), 13566–13574.

14. Radosevich, A. J.; Pu, F.; Chang-Yen, D.; Sawicki, J. W.; Talaty, N. N.; Elsen, N. L.; Williams, J. D.; Pan, J. Y., Ultra-High-Throughput Ambient MS: Direct Analysis at 22 Samples per Second by Infrared Matrix-Assisted Laser Desorption Electrospray Ionization Mass Spectrometry. Anal Chem 2022, 94 (12), 4913–4918.

15. Tu, A.; Said, N.; Muddiman, D. C., Spatially resolved metabolomic characterization of muscle invasive bladder cancer by mass spectrometry imaging. Metabolomics 2021, 17 (8), 70.

16. Bai, H.; Linder, K. E.; Muddiman, D. C., Three-dimensional (3D) imaging of lipids in skin tissues with infrared matrix-assisted laser desorption electrospray ionization (MALDESI) mass spectrometry. Anal Bioanal Chem 2021, 413 (10), 2793–2801.

17. Xu, X.; Meng, Y.; Li, L.; Xu, P.; Wang, J.; Li, Z.; Bian, J., Overview of the Development of Glutaminase Inhibitors: Achievements and Future Directions. J Med Chem 2019, 62 (3), 1096–1115.

18. Gross, M. I.; Demo, S. D.; Dennison, J. B.; Chen, L.; Chernov-Rogan, T.; Goyal, B.; Janes, J. R.; Laidig, G. J.; Lewis, E. R.; Li, J.; Mackinnon, A. L.; Parlati, F.; Rodriguez, M. L.; Shwonek, P. J.; Sjogren, E. B.; Stanton, T. F.; Wang, T.; Yang, J.; Zhao, F.; Bennett, M. K., Antitumor activity of the glutaminase inhibitor CB-839 in triple-negative breast cancer. Mol Cancer Ther 2014, 13 (4), 890–901.

19. Cassago, A.; Ferreira, A. P.; Ferreira, I. M.; Fornezari, C.; Gomes, E. R.; Greene, K. S.; Pereira, H. M.; Garratt, R. C.; Dias, S. M.; Ambrosio, A. L., Mitochondrial localization and structure-based phosphate activation mechanism of Glutaminase C with implications for cancer metabolism. Proc Natl Acad Sci U S A 2012, 109 (4), 1092–7.

20. Palmer, A.; Phapale, P.; Chernyavsky, I.; Lavigne, R.; Fay, D.; Tarasov, A.; Kovalev, V.; Fuchser, J.; Nikolenko, S.; Pineau, C.; Becker, M.; Alexandrov, T., FDR-controlled metabolite annotation for high-resolution imaging mass spectrometry. Nat Methods 2017, 14 (1), 57–60.

