## Supplementary material for "A New Platform for Label-Free, Proximal Cellular Pharmacodynamic Assays: Identification of Glutaminase Inhibitors Using Infrared Matrix-Assisted Laser Desorption Electrospray Ionization Mass Spectrometry": Supportin Information

### Supporting Information

#### Materials

All reagents were purchased from Sigma-Aldrich unless otherwise noted. L-glutamine (1-$^{13}$C) was purchased from Cambridge Isotope Laboratories and 15:0-18:1-d7-PE was purchased from Avanti Polar Lipids. 384-well flat bottom cell culture microtiter plates were purchased from Greiner Bio-One (Item No. 780190), ProxiPlate-384 Plus shallow well plates were purchased from PerkinElmer (Item No. 6008280).

Majority of screening compounds were obtained from AbbVie compound repository. FDA plus library composed of FDA approved compound library and bioactive compounds were purchased from Selleckchem. GAC (72-598) was expressed in *E. Coli* and purified following in-house protocol.

#### IR-MALDESI and MS

We have previously reported the construction of a high throughput IR-MALDESI[1] system used in this work. A 2970 nm IR laser (JGMA Inc.) was focused on the meniscus of liquid samples in the wells and used to generate neutral sample plume traveling upwards and interacting with a constant-on electrospray aligned to the MS inlet. The IR-MALDESI ionization source was coupled to a Q Exactive HF-X mass spectrometer (Thermo Fisher). Automatic gain control was turned off and C-trap injection time was fixed at 20 ms. For cellular metabolite/lipids coverage experiment, MS data were collected in both positive and negative modes, in each mode two mass ranges were scanned at *m/z* 75-300 and *m/z* 300-1200 at a resolving power of 240,000 (FWHM at *m/z* = 200). 80:20 Methanol/water (v/v) with 0.1% formic acid was used as electrospray solvent, L-glutamine (1-$^{13}$C) and 15:0-18:1-d7-PE were included in the electrospray solvent and used as lock masses for both positive and negative mode. For glutaminase cellular and biochemical assay, MS data were collected in negative mode with a mass range of *m/z* 100-200 at a resolving power of 60,000 (FWHM at *m/z* = 200). An extended MS inlet capillary equipped with heating cartridge was necessary to reach the full depth of a standard microtiter plate and aid desolvation. Extended capillary temperature was maintained at 120 °C and MS inlet capillary temperature was set at 400 °C.

Low latency scan mode was used to sync laser firing and MS acquisition, details of this scan mode have been discussed previously[1]. This scan mode provides the fastest speed while maintained the capability of acquiring multiple spectra per sample. Three spectra were taken from each sample and results were averaged during post processing. With experimental conditions described above, a scan speed of 3 minutes per 384-well plate or 2 samples per second was achieved, regardless of biochemical or cellular assay.

#### Cell Culture

A549 (CCL-185™) cell line was obtained from ATCC and cultured at 37°C, 5% $CO_2$ and 90% humidity in T175 flasks (Corning). Cell culture media was made by adding 10% fetal bovine serum (Sigma-Aldrich) and 1% Antibiotic-Antimycotic (100X, Gibco) to DMEM media (Gibco, 11995-040).

#### Compound Dispensing

For cell-based assay, 120 nL of compound were dispensed to 384-well cell culture plates using Echo 655 (Beckman Coulter) to make assay plates. Screening compounds in the main screening library were stored in DMSO stock solutions at 5 mM whereas stock solution concentration was 10 mM for the FDA plus library. The P row (last row on the 384-well plate) was used for controls, consisted of 12 points 1:3 serial dilution of CB839 starting at 10 µM by dispensing 20 nL stock solutions starting at 10 mM, 6 negative control wells (DMSO) and 6 positive control wells (10 µM CB839 in final assay solution). CB839 was used as the positive control compound for both cellular and biochemical assays.

For biochemical assay, compounds were dispensed to assay plates in a similar workflow as the cell-based assay, except 384-well ProxiPlates were used and the dispense volume was reduced to 30 nL per compound. Same control row layout was plated for enzymatic assay although reduced volume at 5 nL was used.

6 point 1:3 serial titration of all available hits from screening were dispensed to make hit confirmation plates with the top assay concentration being 30 µM for both cell-based and enzymatic assay. An additional confirmation with 12 points 1:3 serial titration (top assay concentration was 30 µM) were performed for hits confirmed with 6 points dose responses.

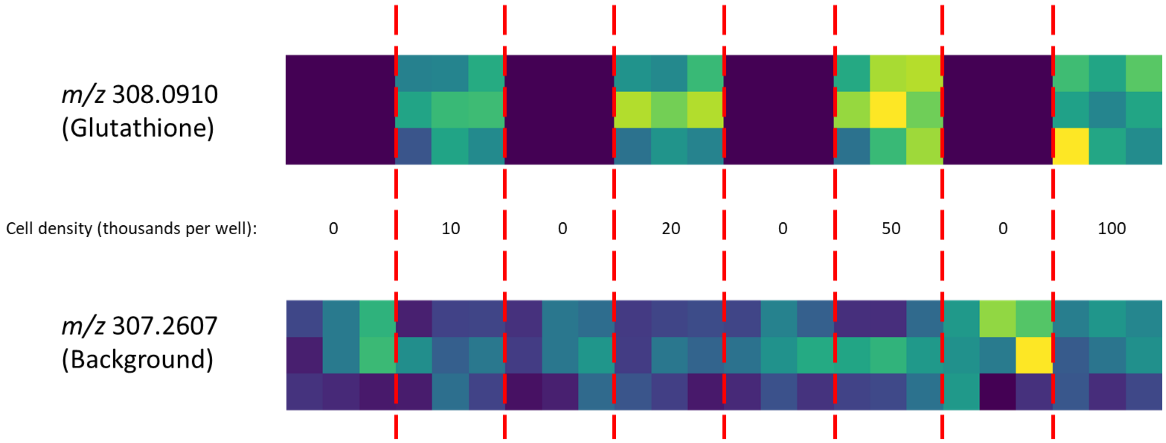


*Figure S 1. Heatmaps of two ions annotated by METASPACE in positive mode. m/z 308.0910 (upper) is likely true signal from the cells because the heatmap pattern agrees with the cell titration pattern. m/z 307.2607 (lower) is likely background signal because it distributes randomly across the wells and was detected in blank wells with no cell.*

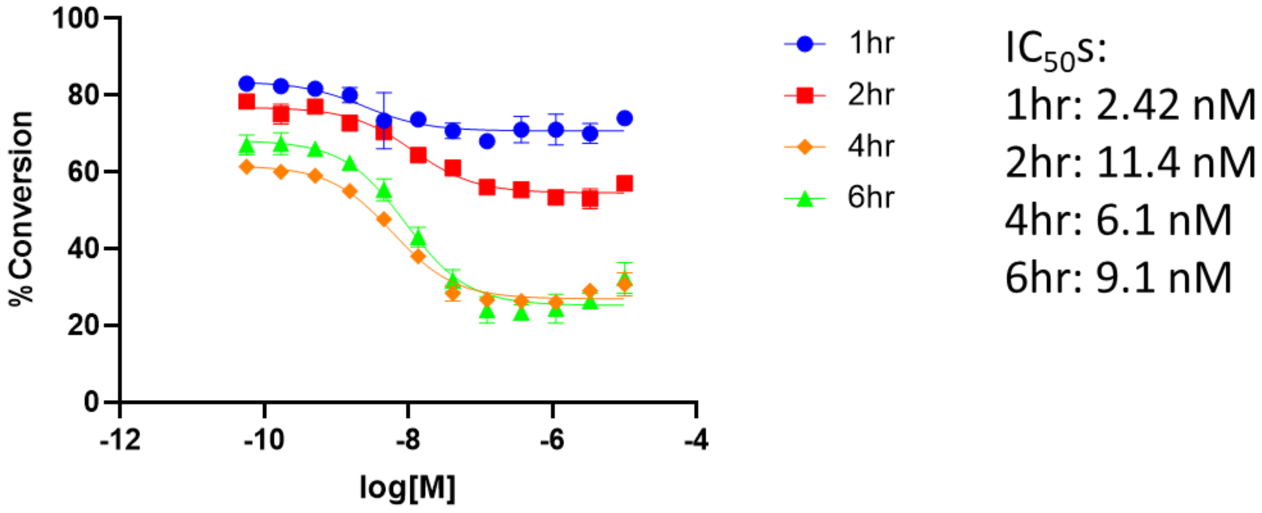


*Figure S 2. Optimization of incubation time for cellular assay. Assay window widens with longer incubation time and stabilizes after 4 hr.*

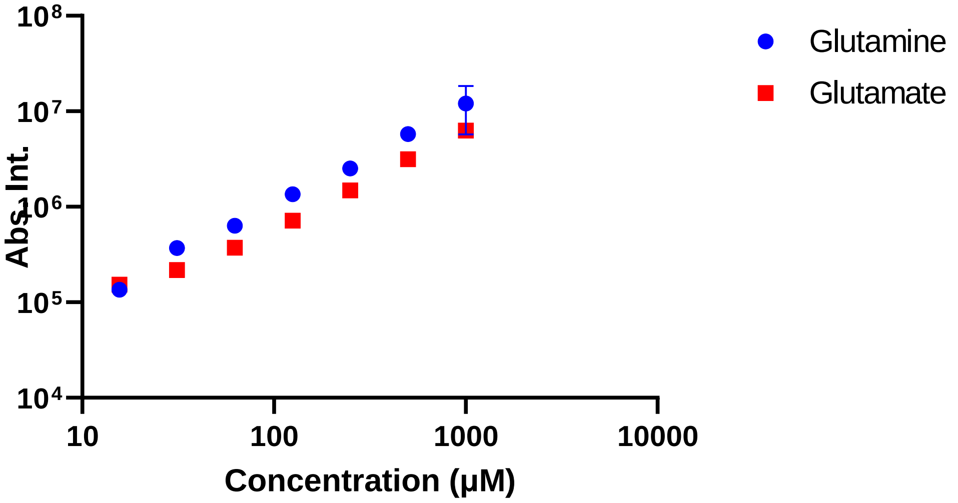


*Figure S 3. Standard curve of glutamine and glutamate in biochemical assay buffer.*

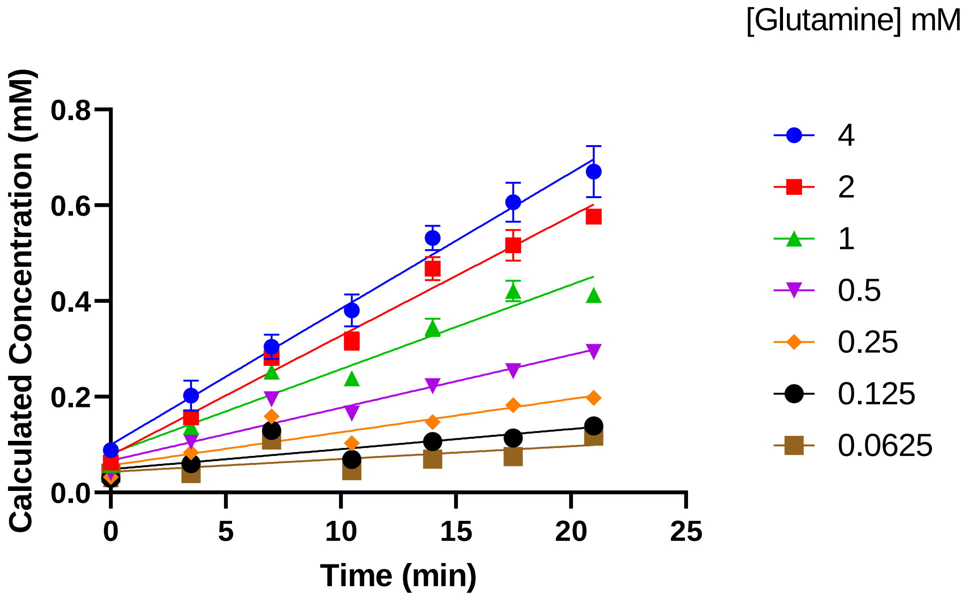


*Figure S 4. Reaction progression curves at different titrated Gln concentrations.*

*Table S 1. Cellular and biochemical assay hit confirmation results.*

| Compound Name | Cellular $IC_{50}$ (μM) | Biochemical $IC_{50}$ (μM) | Dose response Curves (Blue/circle: cellular assay; green/triangle: biochemical assay) |
|---|---|---|---|
| **Crystal Violet** | 0.3788 | 0.5127 | 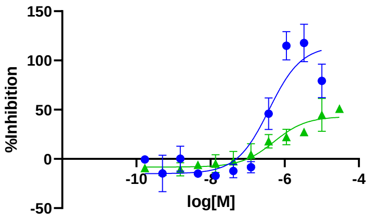 |
| **CB839** | 0.004005 | 0.03858 | 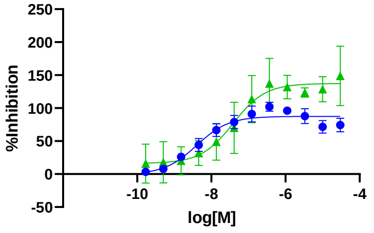 |
| **Sanguinarine chloride** | N/A | 0.3893 | 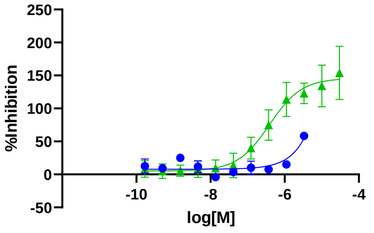 |
| **Compound 1** | 13.83 | 1.761 | 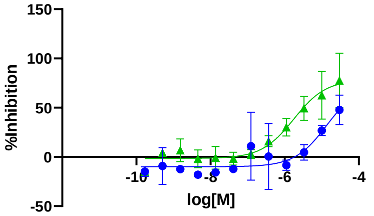 |
| **Compound 2** | 29.18 | 28.18 | 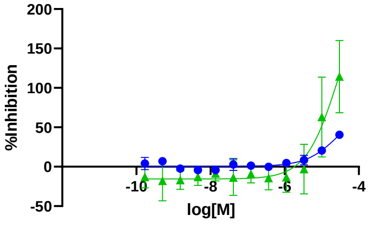 |

| Compound Name | Cellular IC50 (µM) | Biochemical IC50 (µM) | Dose response Curves (Blue/circle: cellular assay; green/triangle: biochemical assay) |
|---|---|---|---|
| **Compound 3** | 18.8 | 7.501 | 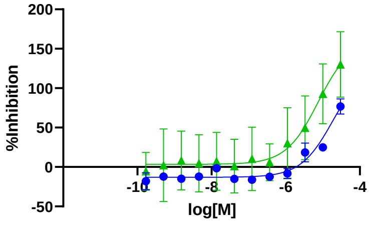 |
| **Compound 4** | 2.712 | 18.11 | 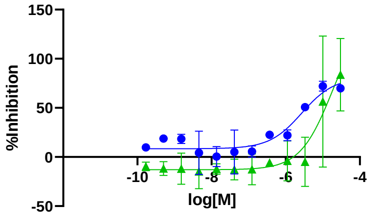 |
| **Compound 5** | 12.8 | 1.149 | 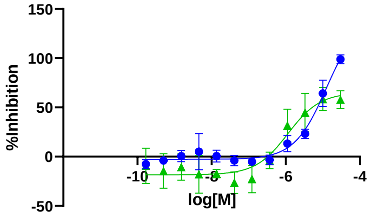 |
| **Compound 6** | 9.011 | 1.618 | 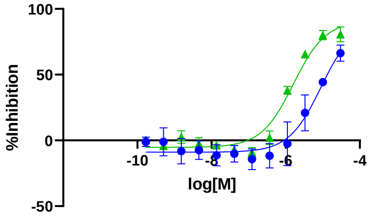 |
| **Compound 7** | 16.44 | 1.192 | 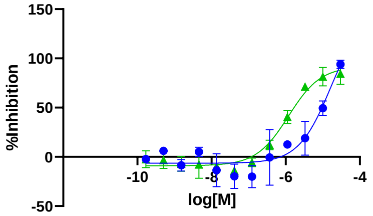 |
| **Compound 8** | 3.069 | 2.397 | 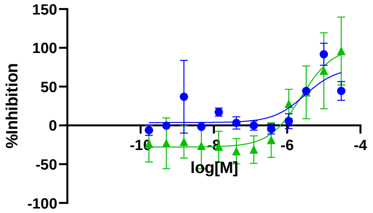 |

| Compound Name | Cellular $IC_{50}$ (µM) | Biochemical $IC_{50}$ (µM) | Dose response Curves (Blue/circle: cellular assay; green/triangle: biochemical assay) |
|---|---|---|---|
| **Compound 9** | 10.61 | 5.924 | 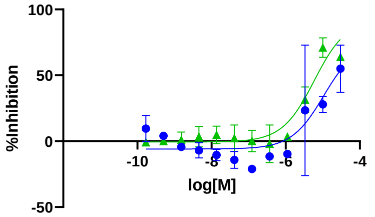  |
| **Compound 10** | 9.659 | 260.9 | 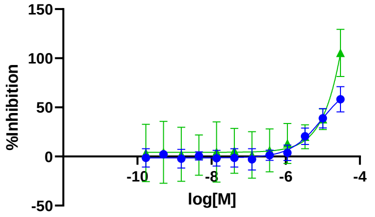  |
| **Compound 11** | 1.568 | 0.2663 | 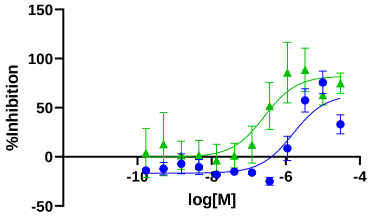  |
| **Compound 12** | 4.437 | 0.4389 | 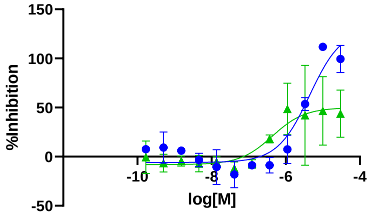  |
| **Compound 13** | 9.535 | N/A | 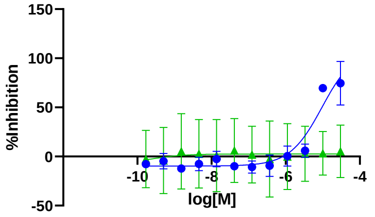  |
| **Compound 14** | 2.321 | N/A | 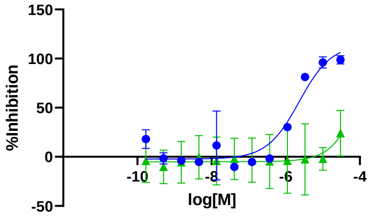  |

| Compound Name | Cellular IC50 (μM) | Biochemical IC50 (μM) | Dose response Curves (Blue/circle: cellular assay; green/triangle: biochemical assay) |
|---|---|---|---|
| **Compound 15** | 1.405 | N/A | 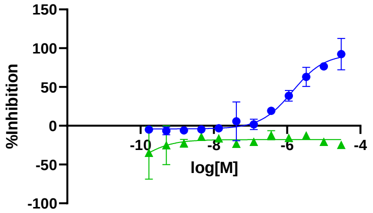 |
| **Compound 16** | 3.594 | N/A | 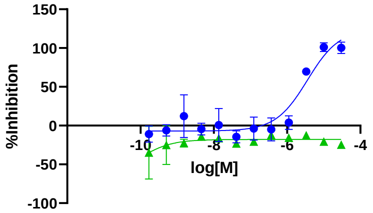 |
| **Compound 17** | 10.44 | N/A | 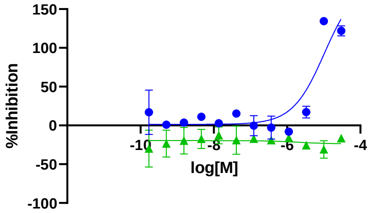 |
| **Compound 18** | 4.489 | N/A | 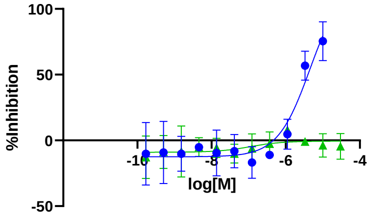 |
| **Compound 19** | 7.886 | N/A | 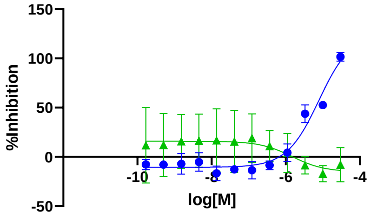 |

### Reference


1. Pu, F.; Radosevich, A. J.; Sawicki, J. W.; Chang-Yen, D.; Talaty, N. N.; Gopalakrishnan, S. M.; Williams, J. D.; Elsen, N. L., High-Throughput Label-Free Biochemical Assays Using Infrared Matrix-Assisted Desorption Electrospray Ionization Mass Spectrometry. *Anal Chem* **2021,** *93* (17), 6792-6800.
